# Abnormal Elevated Connectivity During Language Processing is Associated with Poor Cognitive Performance in Children with Self-limited Epilepsy with Centrotemporal Spikes

**DOI:** 10.1101/2025.06.19.660474

**Authors:** Wendy Qi, Katharine Lee, Kerry C. Nix, Miguel Menchaca, Xiwei She, Lorelei Santa Maria, Wei Wu, Zihuai He, Fiona M. Baumer

## Abstract

Self-Limited Epilepsy with Centrotemporal Spikes (SeLECTS) is associated with language impairments despite seizures originating in the motor cortex, suggesting aberrant cross-network interactions. Here we tested whether functional connectivity in SeLECTS during language tasks predicts language performance. We recorded high-density EEG from right-handed children with SeLECTS (n=31) and age-matched controls (n=32) during verb generation, repetition, and resting tasks. Phonological awareness was assessed using the Comprehensive Test of Phonological Processing-2. Connectivity between bilateral motor cortices and language regions (the left inferior frontal and superior temporal cortices and their right hemisphere homologues) was measured using weighted Phase Lag Index (wPLI). Children with SeLECTS demonstrated significantly elevated connectivity between motor and language regions during language processing. Motor-to-frontal connectivity was higher in SeLECTS during both verb generation and repetition tasks. Frontal-to-temporal connectivity was elevated specifically during verb generation. Higher interhemispheric connectivity (between the left and right hemispheres) during language tasks strongly predicted worse phonological awareness in children with SeLECTS (β= −40 to −61, all p<0.005), but not controls. Together, we found that children with SeLECTS exhibited pathologically elevated connectivity between motor and language networks that was strongly associated with impaired phonological awareness. These findings identify aberrant interhemispheric connectivity as a pathophysiological mechanism underlying language dysfunction and establish EEG-based connectivity measures as a potential biomarker for guiding targeted neuromodulation therapies to treat cognitive impairments in pediatric epilepsy.

## Introduction

Self-limited Epilepsy with Centrotemporal Spikes (SeLECTS) is a common pediatric epilepsy syndrome characterized by frequent spike waves in the motor cortex that are potentiated in sleep^1^. Children with SeLECTS have mild to moderate language impairments^2,3^, often lagging a full standard deviation behind typically developing peers. In addition to causing language dysfunction, SeLECTS affects development of the language network – spatially distant brain regions whose coordinated activity subserves language function^4^. Whereas most children lateralize language to the left hemisphere by age five, children and adolescents with a history of SeLECTS show bilateral temporal and frontal activity during language tasks^5,6^.

Frequent sensorimotor spikes in SeLECTS contribute to language dysfunction, but the mechanism by which they do this is unclear. Spike frequency does not consistently predict language performance^3^. Instead, spikes’ impact on aberrant functional connectivity remains a likely but unproven hypothesis for language dysfunction. Studies using multiple modalities^7,8,9^ find that spikes **acutely** alter functional connectivity between epileptic motor cortices and the language network^10–12^, including the left inferior frontal cortex (IF, Broca’s area), and the left superior temporal (ST, Wernicke’s area). The greater the acute change induced by each spike, the worse the language function^13^, suggesting that susceptibility to spike-induced connectivity disruption is a marker of language vulnerability. Children with SeLECTS also have **chronically** elevated connectivity between spikes, particularly in sleep^12^, but critically, it remains unknown whether the language network itself develops altered connectivity during language processing, independent of acute spike effects, and whether chronic connectivity changes are associated with language function. Elucidating the role of connectivity in language function is clinically relevant, as recent advances in non-invasive neuromodulation techniques, including transcranial magnetic stimulation, have demonstrated the feasibility of selectively modulating cortical connectivity in pediatric populations, opening unprecedented opportunities for targeted cognitive therapeutic interventions in epilepsy^14^.

To test the role of connectivity in language, we compared connectivity in children with and without SeLECTS as they performed three tasks (verb generation, word repetition, and resting state)^15^ during EEG. We then assessed if connectivity was associated with standardized language scores, focusing on phonological awareness as it is consistently reported as abnormal in SeLECTS^3^. We measured connectivity during spike-free trials between six regions: the bilateral motor cortices, left IF and ST language cortices, and right IF and ST homologues using the weighted Phase Lag Index (wPLI), a phase-based connectivity measure robust against volume conduction^16^. We focused on connectivity in the theta (4 to <8 Hz) frequency band, which has been implicated in learning and memory^17,18^. We hypothesized that spikes drive excessive hyperconnectivity between motor and language regions and reduce connectivity between language regions. We expected that high motor-to-language connectivity and low IF to ST connectivity would be associated with worse phonological awareness.

## Methods

### Subjects

Children with a diagnosis of SeLECTS^1^ per ILAE criteria (history of focal seizures during sleep and centrotemporal spikes) were recruited from Lucile Packard Children’s Hospital or surrounding child neurology practices. Typically-developing children with no history of seizures (except simple febrile seizures) or anti-seizure medication (ASM) use were recruited from the community. Inclusion criteria for both groups included being right-handed, 5-13 years old, and born after 35 weeks’ gestation. Children all spoke English though could be bilingual. Exclusion criteria included a history of severe neurologic or systemic problems (i.e. stroke, traumatic brain injury, cardiac or oncologic disease). Children with attention-deficit/hyperactivity disorder (ADHD) were not excluded from either group^19^. Data were collected between February 2021-November 2023. This study was approved by Stanford University’s Institutional Review Board, with informed consent from parents and assent from children.

### Clinical Information

We recorded age, sex, number of lifetime seizures, presence of seizures in sleep only vs. sleep and wakefulness, ASM usage (type), Edinburgh handedness inventory score^20^, bilingual status, socioeconomic status (Hollingshead index score)^21^, and ADHD history.

### Experimental Setup (Figure 1)

Participants sat in front of a computer monitor and speaker in a sound-proof room. High-density EEG was recorded with the Electrical Geodesics NetStation system using 128-channel saline-based caps (sampling rate 1000 Hz) with impedances maintained below 50 kΩ. Stimuli were presented using E-Prime software. Participants completed three tasks (verb generation, repetition, and resting) while staring at a fixation cross to limit eye movement.

**Figure 1.**
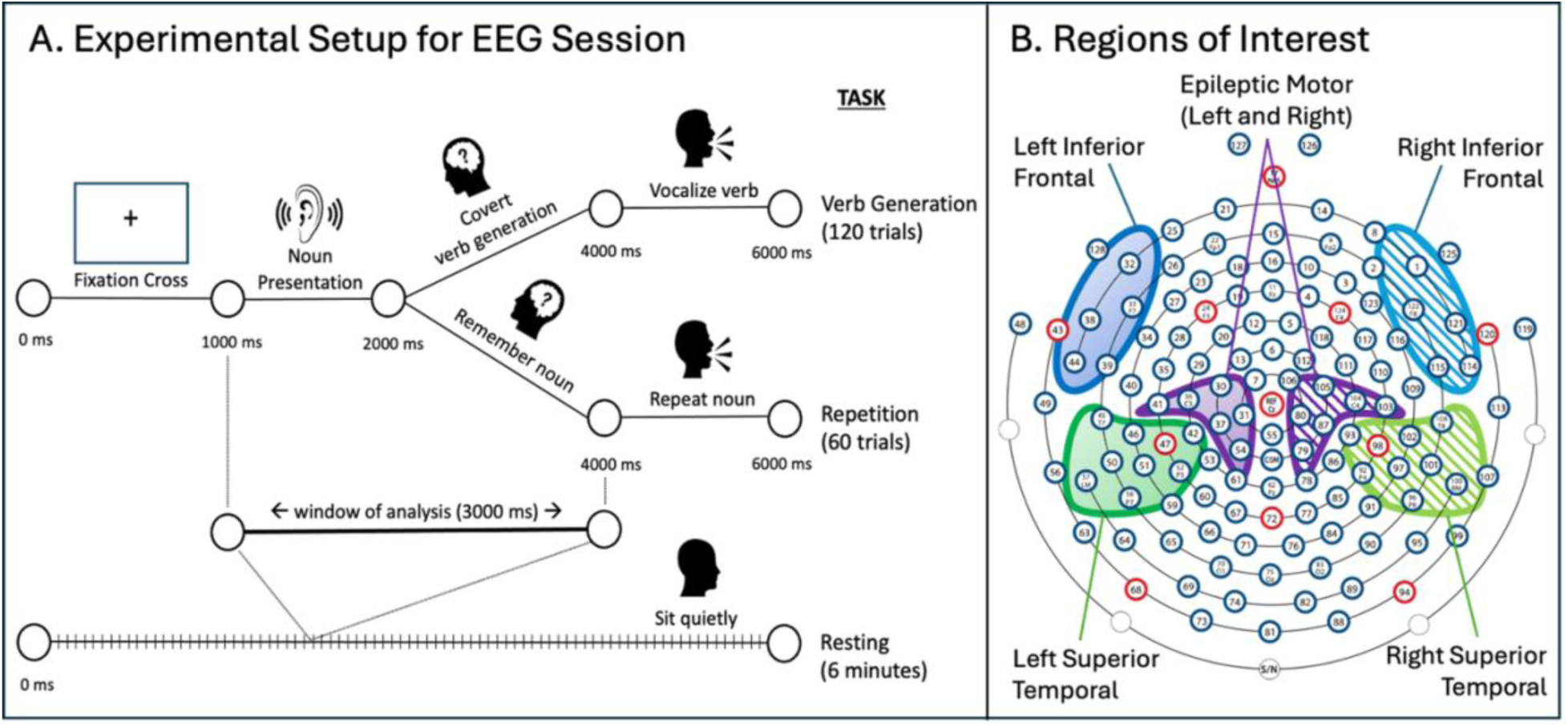
Experimental set-up. (A) Participants completed three tasks (verb generation, repetition, and resting) while high-density EEG was recorded. (B) Topographic plot showing electrodes in each region of interest: inferior frontal (blue), superior temporal (green), and motor (purple) regions with shading indicating hemisphere (solid=left; dashed=right).

The verb generation and repetition tasks began with an auditory presentation of a noun (500-1000 milliseconds in duration) chosen from the Children’s Printed Word Database^22^. Nouns in each block were matched in terms of duration, syllables, and word frequency. The verb generation task – a validated method for activating the language network^23,24^ – consisted of hearing a noun and silently generating an associated “verb or action word.” The repetition task consisted of remembering a noun. After 3 seconds, participants were cued to vocalize the verb or repeat the noun to confirm participation. The repetition task was chosen as a control as it requires working memory and motor planning but lacks the semantic retrieval of verb generation^15^; furthermore, verb generation but not word repetition evokes lateralized EEG changes^15,25^. Two 60-trial blocks of verb generation and one 60-trial block of repetition were conducted. During the resting task, participants sat quietly with their eyes open while looking at the fixation cross for two 3-minute blocks. Block order was randomized on a per participant basis.

### Task Performance

Verb generation responses were coded as correct (appropriate verb); retrieval error (no or non-verb response); or task error (repeated noun). Repetition responses were coded as correct or incorrect.

### Neuropsychological Testing

Testing was performed on a separate day from the EEG by research coordinators (KN, WQ) who had been trained by clinicians. Testing occurred over Zoom given COVID-19 restrictions and was for appropriate scoring as per validated online testing methods^26^. The assessment included the Comprehensive Test of Phonological Processing (CTOPP-2)^27^ Phonological Awareness composite; and the Wechsler Abbreviated Scale Intelligence (WASI-II)^28^ two-subtests form. Phonological awareness was the primary neuropsychological outcome of interest as it is most consistently altered in children with SeLECTS^3,29^.

### Epilepsy Severity

There is not a standardized method for grading epilepsy severity in SeLECTS, but we chose measures based on the GASE survey^30^. We measured the number of lifetime seizures, presence of seizures during sleep only (vs. sleep and wakefulness), presence of seizures lasting for more than 5 minutes, and presence of seizures with secondary generalization, as these factors are important for patient quality of life and could be reliably extracted from medical records. We also categorized patients based on whether spikes were present or absent during the research EEG as this indicates spike activity during wakefulness.

#### Data Analysis

##### Preprocessing

Data were preprocessed in MATLAB (version 2021b) using the Maryland Analysis of Developmental EEG (MADE) pipeline, designed specifically for pediatric EEG preprocessing^31^. For each participant, EEG from all tasks was concatenated, downsampled to 500 Hz, and bandpass filtered (>1Hz and <50Hz). Channels were rejected if values for the Hurst exponent, correlation with other channels, or channel variance exceeded a Z score of three. ICA decomposition was used to identify and remove artifactual ICs^31^. For the verb generation and repetition tasks, three-second epochs were extracted starting at noun onset; resting data was divided into three-second, non-overlapping epochs. Epochs in which the voltage of both ocular electrodes and >10% of non-ocular electrodes exceeded +/-150 μV were rejected. In retained epochs, individual channels exceeding the voltage threshold were interpolated on an epoch-by-epoch basis. Finally, channels rejected earlier in preprocessing were also interpolated. A board-certified epileptologist (FB) manually reviewed remaining epochs and excluded those containing spikes, as spikes acutely increase connectivity^8,12^ and our focus was connectivity elicited by language processing.

Since connectivity estimates are sensitive to epoch number^32^, we aimed to keep trial numbers consistent across subjects and tasks while maximizing data inclusion. Tasks with fewer than 20 epochs after preprocessing^32^ were excluded; thereafter, all epochs up to a maximum of 50 epochs per task were included in connectivity calculations with a random number generator used to reduce to 50 if needed.

##### Extraction of Connectivity Values

Connectivity was calculated using an undirected phase-based measure of brain connectivity, called the wPLI^16^, a robust measure of phase synchronization and connectivity minimally affected by volume conduction^16,32,33^ that is elevated in SeLECTS^12^. We calculated wPLI with cross-spectral density in MATLAB-based Fieldtrip’s connectivity toolbox implementation, as previously published^34^. For each epoch, the phase difference between EEG signals from two electrodes was measured from all time points. The phase differences from all epochs were then averaged to calculate the corresponding wPLI between the electrode pair.

##### Regions of Interest

Regions of interest included the bilateral motor cortices where spikes originate, the left IF and ST regions, and homologous right-sided IF and ST regions^35^. The ST regions also included electrodes overlaying the angular and supramarginal gyri which are important for receptive language^36^. Electrodes corresponding to each region were chosen using standard electrode positions^35^. wPLI values from electrodes spanning regions of interest (e.g. wPLI between left motor and left IF electrodes) were averaged to form a single region-to-region connectivity value. We computed wPLI for twelve region-to-region pairs representing unique connections between bilateral motor, ST and IF regions.

### Statistical Analysis

Analyses were performed with Statistical Analysis System (SAS) OnDemand for Academics^37^. Main analyses compared children with SeLECTS to typically developing controls. Independent t-tests or one-way ANOVA for continuous variables and chi-squared for categorical variables were used to compare groups on: age, sex, handedness, Hollingshead SES Index, IQ, and CTOPP-2 Composite Score. Groups were compared on task performance and EEG data quality metrics (rejected channels, epochs, ICs).

#### Group Differences in Connectivity

Since subjects contributed three connectivity measurements (one per task) to each model, we used a generalized estimating equation (GEE) with an independent correlation matrix^38^ to account for repeated-measures and correlation within individuals. To test if connectivity differed in children with SeLECTS, we fit a model with theta connectivity as the dependent variable and group (SeLECTS/controls), task (verb generation/repetition/resting), and the group by task interaction as independent variables, with adjustment for age and sex; we performed analyses stratified by task. We ran separate models for each of the 12 region-to-region connectivity pairs and considered p < 0.0042 significant as per Bonferroni correction. We hypothesized that children with SeLECTS would have higher connectivity between epileptic motor, IF and ST cortices during all conditions, but lower connectivity between the left IF and ST cortices during the verb generation task. Primary analyses focus on connectivity during the 3000 ms epoch in the theta frequency band (4 to <8 Hz), as theta is related to expressive language processing, word reading, memory, and hippocampal activity^17,18^. Supplementary analyses explored connectivity in: 1) other frequency bands [alpha (8 to 12 Hz), beta (13 to 30 Hz), and gamma (30 to 80 Hz)], also previously reported abnormal in SeLECTS^7,8,12^; and 2) 1000 ms time windows, beginning before noun onset and advancing through the first 2 seconds of the verb generation task.

#### Effect of Antiseizure Medications (ASM) Status on Connectivity

We repeated our primary analysis, this time defining “group” as a 3-level variable (control; children with SeLECTS not taking a daily ASM [SeLECTS-ASM]; and children with SeLECTS taking a daily ASM [SeLECTS+ASM]). We also compared whether epilepsy severity^30^ differed in SeLECTS-ASM and SeLECTS+ASM groups using independent t-tests.

#### Association between Connectivity and Phonological Awareness

We fit a GEE model with CTOPP-2 Composite score as the dependent variable and connectivity, group (SeLECTS/controls), task (verb generation/repetition/resting), and group by task interaction as independent variables, adjusting for age and sex. We conducted analyses stratified by group and task. We hypothesized that children with SeLECTS have different connectivity patterns than controls, and modeled within groups so as not to obscure relevant group-specific relationships. We ran separate models for each region-to-region pair, again using p < 0.0042 as the threshold. We adjusted for unbalanced variables known to be associated with language (SES, IQ, and handedness) in a sensitivity analysis.

## Results

### Subjects

Of the 68 children who consented, one did not participate in the EEG session and four were excluded due to poor data quality, yielding 31 participants with SeLECTS and 32 age and sex-matched controls in the final sample. Three SeLECTS participants and two controls did not complete neuropsychological testing, but their EEG data was included in the connectivity analysis.

### Demographic, Clinical, Neurocognitive, and Data Quality Comparison (Table 1, Supplementary Table 1)

There were no significant group differences in age, sex, or degree of right-handedness. Children with SeLECTS had lower socioeconomic status, IQ, and CTOPP-2 Composite scores than controls. There were no group differences in task performance or EEG data quality measures. Seventeen (55%) SeLECTS participants were taking ASM: 10 levetiracetam, 7 oxcarbazepine.

**Table 1.** Demographics & Cognitive Scores for Children with SeLECTS and Controls. *Edinburgh Handedness Inventory (+1 = strongly right-handed and −1 = left-handed); ADHD: Attention Deficit Hyperactivity Disorder; CTOPP-2: Comprehensive Test of Phonological Processing-2^nd^ Edition; SeLECTS: Self-limited epilepsy with centrotemporal spikes; SES: Socioeconomic Status; WASI-II IQ: Wechsler Abbreviated Scale of Intelligence-2^nd^ Edition, Intelligence Quotient.

| **<u>Demographics</u>** | **<u>Group</u>** |  |  |
| --- | --- | --- | --- |
|  | SeLECTS (n=31) | Controls (n=32) | p-value |
| Age (years) mean, SD | 9.7 +/- 2.0 | 9.1 +/- 2.0 | 0.22 |
| Sex (male), n (%) | 22 (71) | 18 (56) | 0.34 |
| Medication, n (%) | 17 (55) | N/A | N/A |
| Edinburgh Handedness Inventory | 0.82 +/- 0.14 | 0.77 +/- 0.17 | 0.25 |
| Hollingshead SES Index | 54.6 +/- 11.1 | 60.3 +/- 4.4 | 0.01 |
| ADHD status, n (%) | 5 (16) | 6 (19) | 0.73 |

Table 1. Demographics & Cognitive Scores for Children with SeLECTS and Controls.
| **<u>Neuropsychological Assessment</u>** | SeLECTS (n=28) | Controls (n=30) | p-value |
| --- | --- | --- | --- |
| WASI-II IQ | 100.3 +/- 14.2 | 115.2 +/- 14.8 | 0.0004 |
| CTOPP-2 Phonological Awareness Composite Score | 102.3 +/- 14.1 | 113.4 +/- 13.4 | 0.003 |

### Group Differences in Connectivity (Figure 2, Table 2)

Across all regions and conditions, connectivity trended toward higher in the SeLECTS group compared to controls.

**Figure 2.**
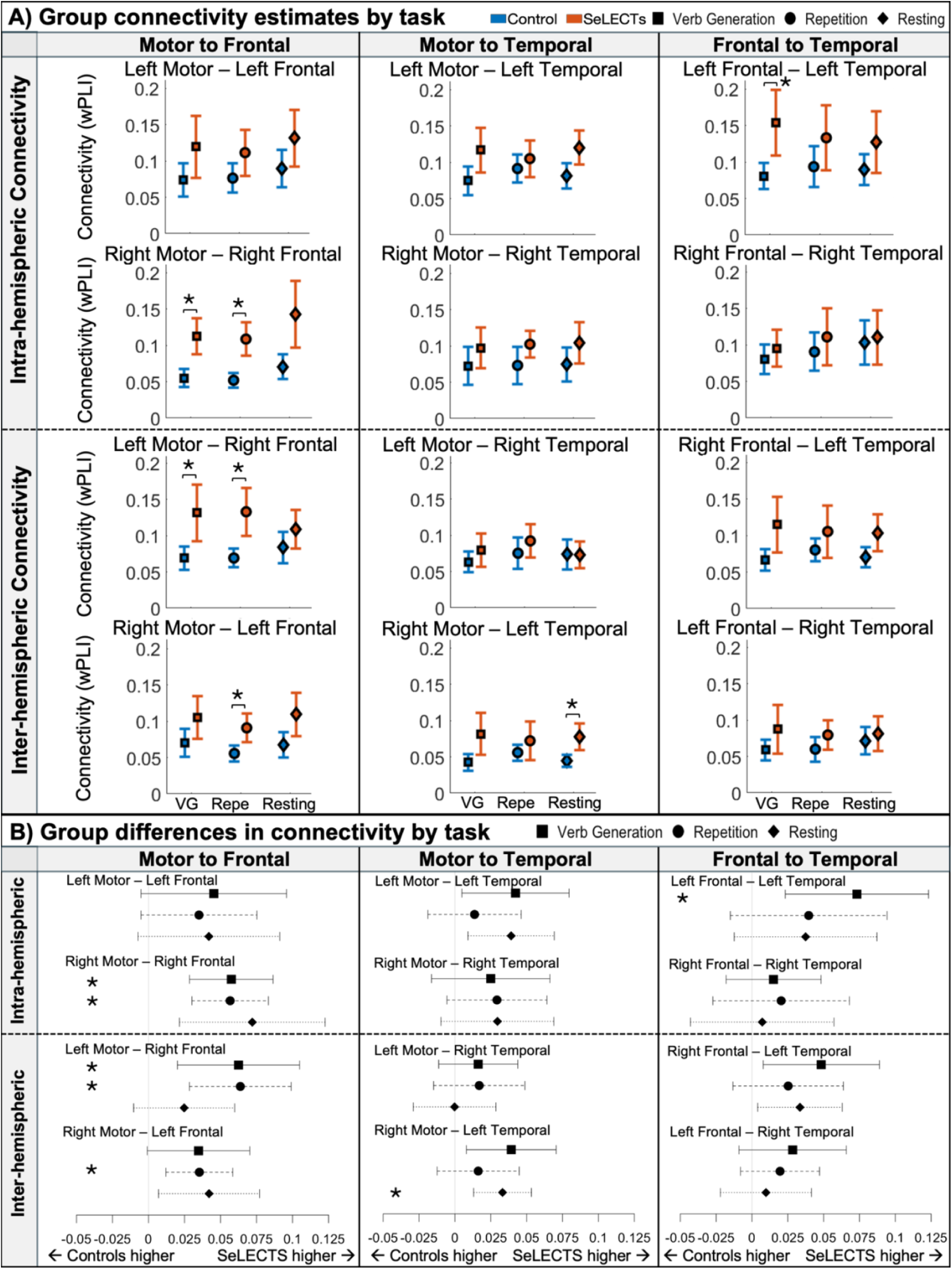
Group differences in theta-band connectivity during all tasks. (A) Box and whisker plots represent estimated marginal mean connectivity with 95% confidence intervals for each group and task. (B) Forest plots represent differences in connectivity estimates between groups with 95% confidence intervals. * indicate p < 0.0042 (adjusted significance threshold). Left column: Motor to Inferior Frontal connectivity; Middle column: Motor to Superior Temporal connectivity; Right column: Inferior Frontal to Superior Temporal connectivity. VG = Verb Generation; Repe = Repetition.

**Table 2.** Group differences in connectivity for Children with SeLECTS and Controls during all tasks. P-values meeting threshold (p<0.0042) are in bold. LFront: Left Inferior Frontal; LTemp: Left Superior Temporal; LMotor: Left Motor; RFront: Right Inferior Frontal; RTemp: Right Superior Temporal; RMotor: Right Motor.

|  |  | <u>Region Pair</u> | <u>Verb Generation</u> |  | <u>Repetition</u> |  | <u>Resting</u> |  |
| --- | --- | --- | --- | --- | --- | --- | --- | --- |
|  |  |  | Estimate<br>(95% CI) | p-value | Estimate<br>(95% CI) | p-value | Estimate<br>(95% CI) | p-value |
| Motor to Frontal | Intra-hemispheric | LMotor — LFront | 0.05<br>(-0.005-0.1) | 0.08 | 0.04<br>(-0.005-0.08) | 0.09 | 0.04<br>(-0.007-0.09) | 0.10 |
|  |  | RMotor — RFront | 0.06<br>(0.03-0.09) | **0.0001** | 0.06<br>(0.03-0.08) | **0.0001** | 0.07<br>(0.02-0.1) | 0.005 |
|  | Inter-hemispheric | LMotor — RFront | 0.06<br>(0.02-0.1) | **0.004** | 0.06<br>(0.03-0.1) | **0.0004** | 0.02<br>(-0.01-0.06) | 0.17 |
|  |  | RMotor — LFront | 0.03<br>(-0.0009-0.07) | 0.06 | 0.04<br>(0.01-0.06) | **0.003** | 0.04<br>(0.007-0.08) | 0.02 |
| Motor to Temporal | Intra-hemispheric | LMotor — LTemp | 0.04<br>(0.005-0.08) | 0.03 | 0.01<br>(-0.02-0.05) | 0.41 | 0.04<br>(0.009-0.07) | 0.01 |
|  |  | RMotor — RTemp | 0.02<br>(-0.02-0.07) | 0.24 | 0.03<br>(-0.006-0.06) | 0.10 | 0.03<br>(-0.01-0.07) | 0.14 |
|  | Inter-hemispheric | LMotor — RTemp | 0.02<br>(-0.01-0.04) | 0.25 | 0.02<br>(-0.01-0.05) | 0.30 | -0.0002<br>(-0.03-0.03) | 0.99 |
|  |  | RMotor — LTemp | 0.04<br>(0.008-0.07) | 0.01 | 0.02<br>(-0.01-0.04) | 0.27 | 0.03<br>(0.01-0.05) | **0.001** |
| Frontal to Temporal | Intra-hemispheric | LFront — LTemp | 0.07<br>(0.02-0.12) | **0.004** | 0.04<br>(-0.02-0.09) | 0.16 | 0.04<br>(-0.01-0.09) | 0.14 |
|  |  | RFront — RTemp | 0.02<br>(-0.02-0.05) | 0.37 | 0.02<br>(-0.03-0.07) | 0.40 | 0.007<br>(-0.04-0.06) | 0.78 |
|  | Inter-hemispheric | RFront — LTemp | 0.05<br>(0.008-0.09) | 0.02 | 0.03<br>(-0.01-0.06) | 0.20 | 0.03<br>(0.004-0.06) | 0.03 |
|  |  | LFront — RTemp | 0.03<br>(-0.009-0.07) | 0.14 | 0.02<br>(-0.008-0.05) | 0.16 | 0.01<br>(-0.02-0.04) | 0.55 |

#### Motor to Inferior Frontal Connectivity

SeLECTS children had significantly higher connectivity between the right IF region and both the left and right motor regions during the verb generation and repetition tasks. Connectivity was elevated between the left IF and right motor regions only during repetition.

#### Motor to Superior Temporal Connectivity

Connectivity was elevated in SeLECTS between the left ST and right motor regions during resting.

#### Inferior Frontal to Superior Temporal Connectivity

Connectivity between the left IF and left ST regions – the regions most implicated in language processing – was elevated in SeLECTS specifically during the verb generation task.

### Connectivity in other Frequency Bands (Supplementary Tables 2-4)

There were no significant group differences in the alpha, beta, or gamma bands.

### Connectivity over Specific Time Windows (Supplementary Figure 1)

Baseline connectivity before noun onset did not differ between groups. The SeLECTS group had higher connectivity than controls between the bilateral motor and bilateral IF and between bilateral motor and left ST, most prominently from 0.5 to 1.5 seconds after noun onset. Left IF to left ST connectivity was also elevated in SeLECTS in this period. Results were consistent with analyses using longer 3 second epochs.

### Effect of Antiseizure Medications on Connectivity (Figure 3, Supplementary Tables 5-9)

When comparing three groups (SeLECTS+ASM, SeLECTS-ASM, and controls), there were significant group differences in SES, IQ, and CTOPP-2 scores. The same region pairs as described above showed significantly higher connectivity in the SeLECTS+ASM group vs. controls. Connectivity in the SeLECTS-ASM group did not differ significantly from that in controls, but notably their connectivity values often fell in between those of controls and SeLECTS+ASM. The SeLECTS+ASM group had higher connectivity than the SeLECTS-ASM group in two regions (left IF to left ST regions during verb generation; left IF to right ST during repetition). Epilepsy severity did not differ between the SeLECTS+ASM and SeLECTS-ASM groups.

**Figure 3.**
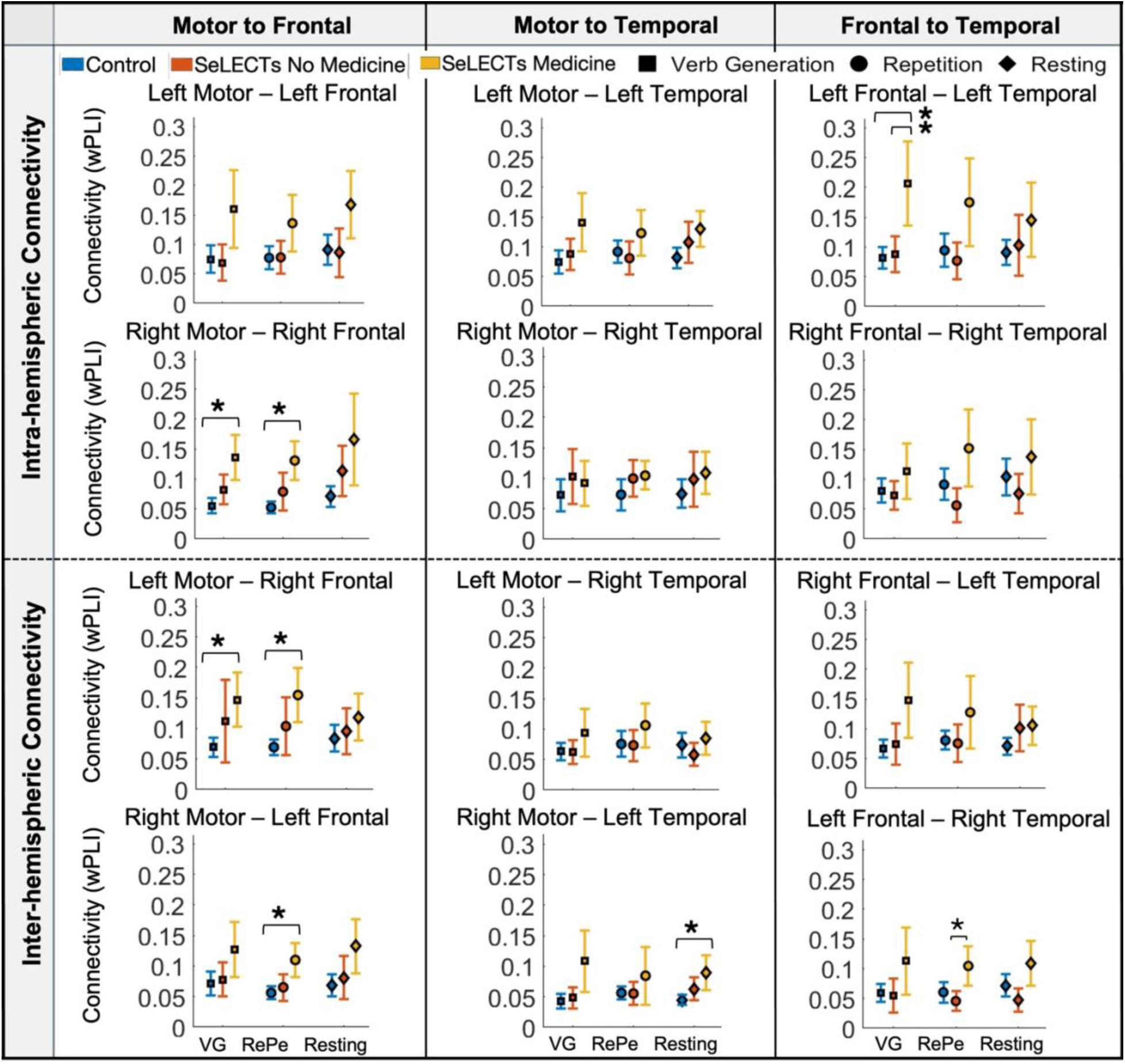
Impact of Antiseizure Medication (ASM) Use on Connectivity. Plots represent estimated marginal mean connectivity with 95% confidence interval for each group and task. * indicate p < 0.0042 (adjusted significance threshold). Left column: Motor to Inferior Frontal connectivity; Middle column: Motor to Superior Temporal connectivity; Right column: Inferior Frontal to Superior Temporal connectivity. VG = Verb Generation; Repe = Repetition.

### Association between Connectivity and Phonological Awareness (Figure 4, Tables 3-4; Supplementary Figures 2-3 & Tables 10-11)

#### SeLECTS

High *interhemispheric* connectivity (connectivity between regions in opposite hemispheres) during the verb generation task, and to a lesser extent during the repetition task, were strongly associated with poor phonological awareness. There were significant associations in five of the six interhemispheric region-pairs during the verb generation task and three region-pairs during the repetition task. *Intra-hemispheric connectivity* (connectivity between regions in the same hemisphere) was not associated with phonological awareness. Resting connectivity was not associated with phonological awareness in SeLECTS. Associations remained significant in four of six interhemispheric pairs even after adjusting for SES, IQ, and handedness.

#### Controls

Higher right motor to right ST connectivity in resting was associated with lower phonological awareness. There were no other significant associations in controls. Associations were significant in three interhemispheric pairs after adjusting for SES, IQ, and handedness.

**Figure 4.**
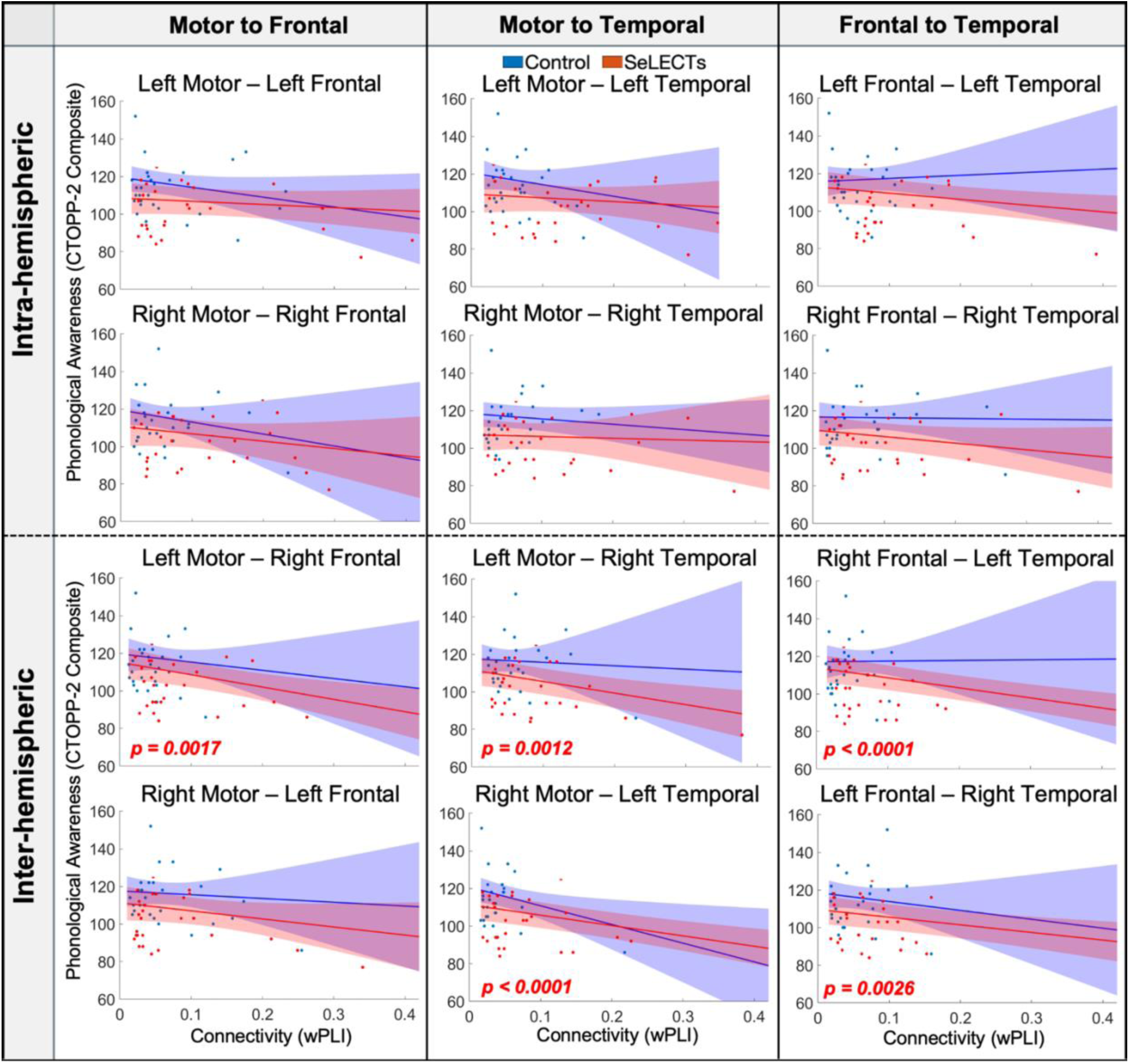
Association between Connectivity and Clinical Language Performance. Scatter plots of mean connectivity vs. CTOPP-2 Composite scores for each subject and estimated marginal fit lines with 95% confidence intervals. P-values meeting threshold (p<0.0042) noted and color coded to group (red=SeLECTS; blue=controls). Left column: Motor to Inferior Frontal connectivity; Middle column: Motor to Superior Temporal connectivity; Right column: Inferior Frontal to Superior Temporal connectivity.

**Table 3.**
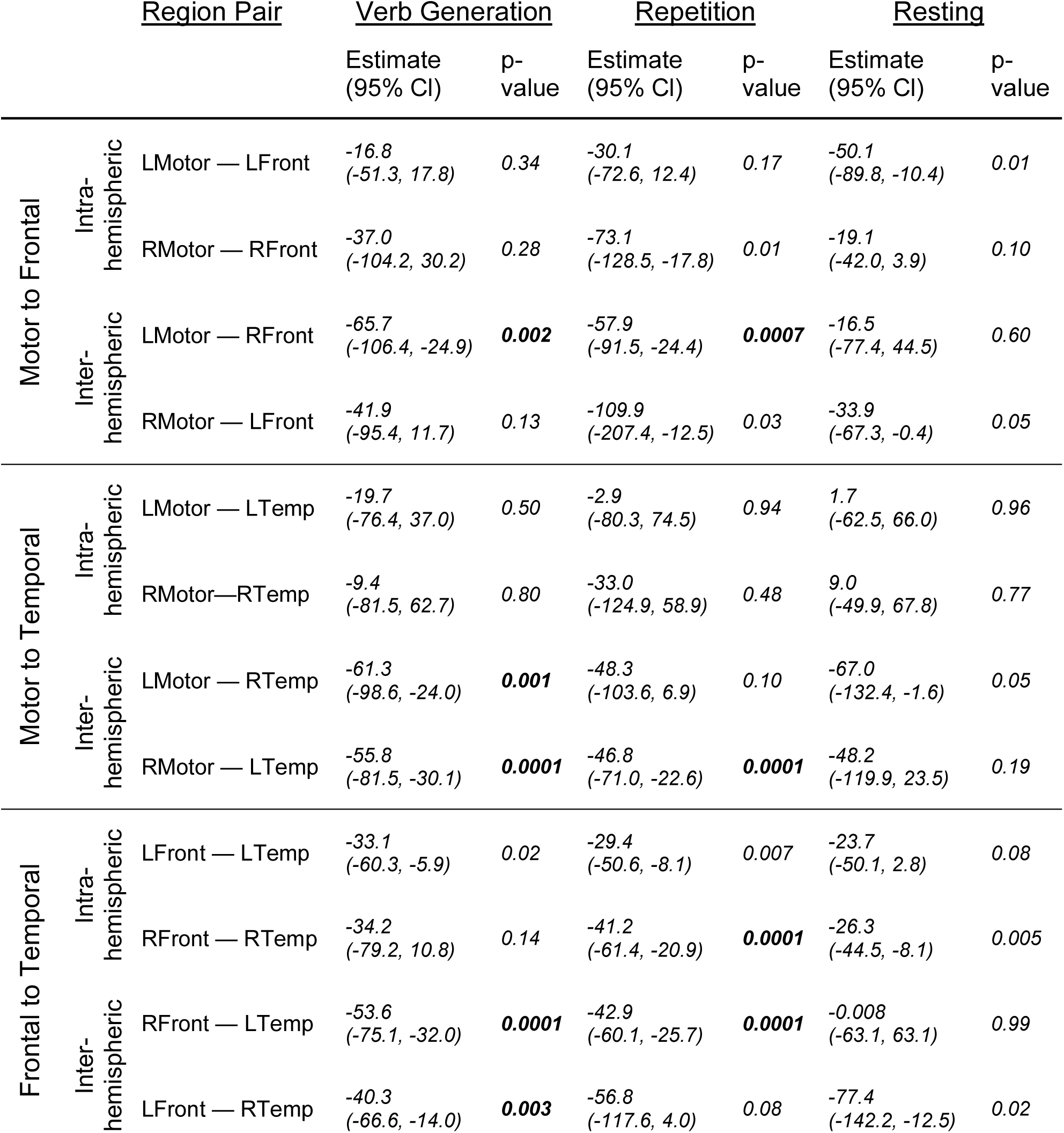
Association between Clinical Language Performance and Connectivity in Children with SeLECTS. P-values meeting threshold (p<0.0042) are in bold. LFront: Left Inferior Frontal; LTemp: Left Superior Temporal; LMotor: Left Motor; RFront: Right Inferior Frontal; RTemp: Right Superior Temporal; RMotor: Right Motor.

**Table 4.** Association between Clinical Language Performance and Connectivity in Controls. P-values meeting threshold (p<0.0042) are in bold. LFront: Left Inferior Frontal; LTemp: Left Superior Temporal; LMotor: Left Motor; RFront: Right Inferior Frontal; RTemp: Right Superior Temporal; RMotor: Right Motor.

|  |  | <u>Region Pair</u> | <u>Verb Generation</u> |  | <u>Repetition</u> |  | <u>Resting</u> |  |
| --- | --- | --- | --- | --- | --- | --- | --- | --- |
|  |  |  | Estimate<br>(95% CI) | p-value | Estimate<br>(95% CI) | p-value | Estimate<br>(95% CI) | p-value |
| Motor to Frontal | Intra-hemispheric | LMotor — LFront | -52.5<br>(-119.4, 14.3) | 0.12 | -29.3<br>(-102.6, 44.1) | 0.43 | -44.3<br>(-103.8, 15.3) | 0.15 |
|  |  | RMotor — RFront | -60.8<br>(-173.7, 52.1) | 0.29 | -95.9<br>(-252.2, 60.4) | 0.23 | -76.9<br>(-163.3, 9.5) | 0.08 |
|  | Inter-hemispheric | LMotor — RFront | -42.2<br>(-145.7, 61.3) | 0.42 | -31.7<br>(-137.6, 74.2) | 0.56 | -15.8<br>(-77.9, 46.3) | 0.62 |
|  |  | RMotor — LFront | -19.5<br>(-114.6, 75.6) | 0.69 | -128.5<br>(-233.0, -23.9) | 0.02 | -80.9<br>(-163.4, 1.5) | 0.05 |
| Motor to Temporal | Intra-hemispheric | LMotor — LTemp | -60.5<br>(-183.6, 62.6) | 0.34 | -61.6<br>(-132.6, 9.4) | 0.09 | 39.9<br>(-23.3, 103.2) | 0.22 |
|  |  | RMotor — RTemp | -28.1<br>(-80.0, 23.9) | 0.29 | -31.2<br>(-87.6, 25.1) | 0.28 | -58.2<br>(-87.5, -28.9) | **0.0001** |
|  | Inter-hemispheric | LMotor — RTemp | -17.9<br>(-164.7, 129.0) | 0.81 | -66.6<br>(-113.9, -19.4) | 0.006 | -60.9<br>(-122.6, 0.8) | 0.05 |
|  |  | RMotor — LTemp | -98.2<br>(-179.4, -17.0) | 0.02 | -135.2<br>(-242.1, -28.3) | 0.01 | -18.8<br>(-232.7, 195.0) | 0.86 |
| Frontal to Temporal | Intra-hemispheric | LFront — LTemp | 18.2<br>(-78.7, 115.2) | 0.71 | 2.3<br>(-41.1, 45.6) | 0.92 | 10.2<br>(-49.5, 69.9) | 0.74 |
|  |  | RFront — RTemp | -1.5<br>(-83.2, 80.1) | 0.97 | -47.5<br>(-95.3, 0.32) | 0.05 | -51.8<br>(-89.0, -14.7) | 0.006 |
|  | Inter-hemispheric | RFront — LTemp | 4.1<br>(-125.0, 133.1) | 0.95 | -12.3<br>(-90.6, 65.9) | 0.76 | 16.8<br>(-59.7, 93.2) | 0.67 |
|  |  | LFront — RTemp | -47.2<br>(-143.9, 49.4) | 0.34 | -93.2<br>(-162.2, -24.2) | 0.008 | -76.2<br>(-142.9, -9.4) | 0.03 |

## Discussion

We investigated whether language problems associated with epilepsy disrupt functional connectivity. Using high-density EEG recordings, we measured connectivity between the motor cortices – the source of interictal spikes and seizures in SeLECTS – and IF and ST language regions. We found three key patterns. First, while performing a language task, children with SeLECTS have significantly higher connectivity between motor and IF regions than controls, with the largest differences involving the *right* IF region. Second, contrary to our hypothesis, connectivity between IF and ST language regions in the left hemisphere was also *elevated* in SeLECTS. Third, higher interhemispheric connectivity was strongly associated with worse phonological awareness in children with SeLECTS, suggesting that excessive connectivity impedes language processing. Our findings suggest hyperconnectivity contributes to language difficulties in SeLECTS, offering insights into therapeutic targets.

### Motor to Inferior Frontal (IF) Connectivity Differences

Motor-to-IF connectivity is elevated in SeLECTS specifically during language tasks but not during resting. We were surprised that differences were task specific as we hypothesized that motor cortex hyperconnectivity would be driven by spikes and hence present regardless of cognitive demands. Given that motor-to-IF hyperconnectivity is observed during both verb generation and repetition tasks, we speculate it is elicited by a shared cognitive component – such as attention, working memory or articulatory planning^15^ – rather than semantic retrieval component unique to verb generation. The predominance of right IF findings is particularly notable, as this region has been implicated in inhibitory control^39^, a key component for attentional processes which are disrupted in SeLECTS^40^. Supporting this, a spike-timed EEG-fMRI study^41^ found that centrotemporal spikes activate both the right IF and the caudate nucleus, structures important for attention regulation. Alternatively, the right IF region may play a more prominent role in expressive language in SeLECTS. Verb generation elicits excessive right frontal activation both in children with active SeLECTS^2,42^ and in adolescents with a history of the disorder^43^. Lateralization of the anterior language network may be specifically disrupted by SeLECTS pathology^42^, requiring affected children to recruit additional right hemispheric regions to maintain normal task performance.

### Motor to Superior Temporal (ST) Connectivity Differences

Motor-to-ST connectivity was similar between groups, with the only difference being elevated right motor to left ST connectivity during resting but not during tasks. Prior studies have found that SeLECTS disrupts the default mode network (DMN), a set of interconnected brain regions that coactivate during the resting state. DMN disruption affects cognitive efficiency by interfering with resource allocation between task-positive and task-negative networks, contributing to language difficulties^44^. Resting connectivity and DMN dysregulation may represent another mechanism through which motor-cortex spikes impact distributed cognitive networks, including those supporting language processing.

### Frontal to Temporal Connectivity Differences

Contrary to our hypothesis and previous neuroimaging studies showing reduced structura^l29^ and resting-state functiona^l45^ connectivity in the left hemisphere language network, we found *increased* left IF to left ST connectivity in SeLECTS specifically during the verb generation task. This suggests that language network connectivity during semantic retrieval is affected in SeLECTS. One explanation is that elevated connectivity reflects delayed maturation of the language network. This interpretation aligns with fMRI studies showing that activation in higher-level language regions decreases with age as language processing becomes more efficient^46^ and increases again (even in adulthood) when tasks become more semantically challenging^47^. Children with SeLECTS may require more intensive utilization of language regions to achieve the same behavioral output as typically developing children, resulting in higher measured connectivity. Supporting this interpretation, task performance did not differ between groups despite these neurophysiological differences, suggesting compensatory mechanisms. A second interpretation is that higher connectivity represents accelerated language network development; one MEG study found that theta band connectivity increases over time in children and is associated with improving language function^17^. We think this is unlikely in SeLECTS, where higher connectivity is associated with poorer phonological awareness. A third explanation is that SeLECTS induces atypical development of the language network, much as seizures and spikes can alter connectivity in animal models. Adolescents with a history of resolved SeLECTS still have different EEG activity during verb generation, indicating lasting network reorganization^43^. The critical distinction between delayed versus atypical development has significant implications for prognosis and intervention. With delayed development, connectivity abnormalities may normalize with age or seizure resolution whereas atypical development demands targeted early interventions. Longitudinal studies examining how connectivity patterns evolve during active epilepsy and after resolution, and the relationship between connectivity and language, will be crucial to address this question.

### Effect of Antiseizure Medications (ASMs)

ASM effects vary by region. Motor-to-IF connectivity follows a gradient: controls have lowest connectivity, unmedicated children with SeLECTS have mid-level connectivity, and medicated children with SeLECTS have highest connectivity. Connectivity in unmedicated children with SeLECTS more closely resembles that of controls than that of medicated children, but small sample size limits power. Importantly, left IF-to-ST connectivity is significantly higher in children taking ASMs than either controls or unmedicated children. The easiest explanation is that more severe epilepsy is associated with both higher connectivity and ASM prescription; children on ASMs had more lifetime seizures thus higher connectivity. An alternative explanation is that ASMs directly increase connectivity, but prior literature is mixed. When followed over two years, EEG functional connectivity remained stable in children with SeLECTS who started an ASM but increased in those who remained unmedicated^48^. A MEG study found an increase in functional connectivity in the default mode network after initiation of ASM^49^. Both studies support ASMs’ ability to alter connectivity in SeLECTS but an understanding of how and whether they normalize connectivity is limited given the lack of a control group. A cross-sectional fMRI study also found that both medicated and unmedicated children with SeLECTS have different functional connectivity than controls, but it was *reduced* in children on ASMs and *elevated* in children off ASMs^50^. Disentangling the effects of epilepsy severity vs. ASMs on connectivity is important as there is still equipoise regarding the decision to start ASMs.

### Association between Connectivity and Phonological Awareness

Higher interhemispheric connectivity during the verb generation and repetition tasks is associated with worse phonological awareness in children with SeLECTS and not controls. Two fMRI studies also found that interhemispheric connectivity is associated with language abilities in SeLECTS. One highly concordant study^13^ noted that higher motor-to-left-temporal connectivity was associated with lower verbal and full-scale IQ in SeLECTS. In contrast, the second study found that higher left motor to right frontal connectivity was associated with *better* language scores^2^. While the direction of the association varies, both suggest that interhemispheric connectivity plays a role in language function. Notably, both studies looked at connectivity during the resting state, whereas we focused on a language task. Children with dyslexia (which also impairs phonological awareness)^51^ also have greater interhemispheric connectivity than controls during a word-tracking task and high connectivity is associated with poor phonological awareness. We propose that high interhemispheric connectivity is associated with poor language performance because it signifies that a segregated left-lateralized language network has not (yet) developed. Corroborating this, appropriate activation of limited, task-appropriate brain networks (rather than wider brain regions) is associated with better cognitive performance in SeLECTS^44,52^ and other focal epilepsies^53^. Whether excessive connectivity is a pathology that directly interferes with language function vs. a compensatory response to underlying network dysfunction could be tested using non-invasive neurostimulation.

#### Limitations

First, while the group connectivity differences were pronounced, clear task-specific differences within groups were minimal, suggesting that the connectivity measure was not sensitive to differences in verb generation and repetition; future studies can consider nonword repetition tasks, which uncouples phonological awareness from vocabulary knowledge and may better differentiate language-specific neural processes. Notably, group connectivity differences became more apparent with the increasing cognitive load required for verb generation. Second, our tasks did not capture natural language complexity, and future studies measuring naturalistic language abilities would enhance generalizability. Third, averaging connectivity over a three-second window may obscure rapid language-related changes. Our analyses using a 1-second window supported our primary findings, but more temporally precise analyses might be beneficial. Fourth, our scalp-based connectivity measurements lack spatial specificity. While source localization would have improved anatomical precision, this approach was difficult due to the lack of individual neuroimaging data. Nevertheless, we observed clear regional differences in connectivity that varied with task demands in sensor space, suggesting that our approach captured meaningful functional distinctions between regions. Fifth, our groups were not perfectly matched for all variables that may affect language (e.g. socioeconomic status; IQ)^54^. Our supplementary analyses adjusting for these variables did not alter the negative relationship between high connectivity and phonological awareness, but future studies with better matching would strengthen conclusions about epilepsy-specific effects. Finally, different ASMs may have different effects on connectivity, and we lack the power to reliably explore these differences.

#### Conclusion

Children with SeLECTS have aberrant hyperconnectivity between epileptic motor and language cortices that arises specifically during language processing and is strongly associated with phonological processing deficits. Our study provides support for suppressing hyperconnected motor-language pathways, and particularly interhemispheric connections, to normalize network function and improve language outcomes in children with SeLECTS, establishing a foundation for precision neuromodulation approaches in pediatric epilepsy.

## Acknowledgements

Funding was provided by K23 NS116110, the Doris Duke & Rita Allen Foundations, a gift from the Principe & O’Farrell family, and the Wu Tsai Neurosciences Institute to FMB. Funding was provided by the Maternal & Child Health Research Institute to XS. We also thank Dr. Baldeweg for sharing the word lists used in this study.

## Author Contributions

W.Q.: conceptualization, data curation, formal analysis, software, writing – original draft, writing – review & editing. K.L.: conceptualization, data curation, formal analysis, software, writing – original draft, writing – review & editing. K.C.N.: methodology, software, data curation, validation, writing – review & editing. M.M.: data curation, validation, writing – review & editing. X.S.: software, validation, writing – review & editing. L.S.M.: formal analysis, writing – review & editing. W.W.: methodology, software, writing – review & editing. Z.H.: methodology, writing – review & editing. F.M.B.: conceptualization, methodology, software, data curation, formal analysis, writing – original draft, writing – review & editing, project administration, funding acquisition.

## Potential Conflicts of Interest

The authors declare no conflicts of interest.

## Data Availability

The datasets generalized and analyzed in the study are not currently publicly available due to privacy concerns with de-identification of clinical data. However, the full dataset is available from the corresponding author by request.

## Notes

### Competing Interest Statement

The authors have declared no competing interest.

