## Supplementary Tables and Figures for "Abnormal Elevated Connectivity During Language Processing is Associated with Poor Cognitive Performance in Children with Self-limited Epilepsy with Centrotemporal Spikes"

**Supplemental Table 1.** Preprocessing and task performance results for children with SeLECTS and controls.

| **Data Quality** | **Group** | | | |
|---|---|---|---|---|
| | SeLECTS (n=31) | Controls (n=32) | t-statistic | p-value |
| Number of rejected channels | *5.6 +/- 2.0* | *5.4 +/- 2.2* | *0.38* | *0.70* |
| % of rejected ICs | *23.4% +/- 11.1%* | *20.4% +/- 5.2%* | *1.4* | *0.18* |
| **Number of Epochs** | | | | |
| Verb Generation Task | *47.5 +/- 6.3* | *49.3 +/- 2.0* | *1.6* | *0.12* |
| Repetition Task | *40.4 +/- 9.3* | *43.0 +/- 7.9* | *1.2* | *0.24* |
| Resting Task | *50 +/- 0* | *49.7 +/- 1.6* | *1.1* | *0.28* |
| **Task Response** | | | | |
| Verb Response | *87.4 +/- 26.9* | *88.6 +/- 33.7* | *0.15* | *0.88* |
| Task Error | *3.2 +/- 9.0* | *1.1 +/- 1.5* | *1.2* | *0.24* |
| Retrieval Error | *23.5 +/- 17.8* | *32.3 +/- 22.7* | *1.7* | *0.1* |

*ICs = Independent components

**Supplemental Table 2.** Group differences in connectivity for Children with SeLECTS in ALPHA band during all tasks.

| | | Region Pair | Verb Generation | | Repetition | | Resting | |
|---|---|---|---|---|---|---|---|---|
| | | | Estimate (95% CI) | p-value | Estimate (95% CI) | p-value | Estimate (95% CI) | p-value |
| Motor to Frontal | Intra-hemispheric | LMotor — LFront | *0.02 (-0.03, 0.07)* | *0.44* | *0.04 (-0.01, 0.09)* | *0.13* | *0.05 (-0.02, 0.1)* | *0.16* |
| | | RMotor — RFront | *0.006 (-0.04, 0.05)* | *0.78* | *0.04 (-0.02, 0.09)* | *0.20* | *0.06 (-0.009, 0.1)* | *0.09* |
| | Inter-hemispheric | LMotor — RFront | *0.02 (-0.03, 0.07)* | *0.51* | *0.05 (-0.02, 0.1)* | *0.19* | *0.003 (-0.07, 0.08)* | *0.94* |
| | | RMotor — LFront | *0.03 (-0.01, 0.07)* | *0.14* | *0.03 (-0.02, 0.08)* | *0.18* | *0.06 (-0.005, 0.1)* | *0.07* |
| Motor to Temporal | Intra-hemispheric | LMotor — LTemp | *0.07 (-0.001, 0.1)* | *0.05* | *0.03 (-0.04, 0.09)* | *0.41* | *0.04 (-0.01, 0.1)* | *0.13* |
| | | RMotor—RTemp | *0.007 (-0.05, 0.06)* | *0.80* | *-0.001 (-0.05, 0.05)* | *0.97* | *0.05 (-0.001, 0.1)* | *0.05* |
| | Inter-hemispheric | LMotor — RTemp | *0.02 (-0.03, 0.06)* | *0.47* | *0.03 (-0.02, 0.07)* | *0.25* | *0.006 (-0.03, 0.05)* | *0.77* |
| | | RMotor — LTemp | *0.07 (0.01, 0.1)* | *0.01* | *0.04 (-0.01, 0.09)* | *0.15* | *0.04 (-0.001, 0.08)* | *0.06* |
| Frontal to Temporal | Intra-hemispheric | LFront — LTemp | *0.08 (0.01, 0.1)* | *0.02* | *0.05 (-0.02, 0.1)* | *0.14* | *-0.03 (-0.09, 0.04)* | *0.38* |
| | | RFront — RTemp | *-0.03 (-0.1, 0.04)* | *0.42* | *-0.02 (-0.1, 0.06)* | *0.63* | *-0.01 (-0.09, 0.06)* | *0.69* |
| | Inter-hemispheric | RFront — LTemp | *0.06 (-0.001, 0.1)* | *0.05* | *0.06 (-0.006, 0.1)* | *0.08* | *-0.01 (-0.07, 0.05)* | *0.71* |
| | | LFront — RTemp | *-0.02 (-0.08, 0.04)* | *0.50* | *-0.009 (-0.07, 0.05)* | *0.79* | *0.01 (-0.05, 0.08)* | *0.66* |

P-values meeting threshold ($p<0.0042$) are in bold. LFront: Left Inferior Frontal; LTemp: Left Superior Temporal; LMotor: Left Motor; RFront: Right Inferior Frontal; RTemp: Right Superior Temporal; RMotor: Right Motor.

**Supplemental Table 3.** Group differences in connectivity for Children with SeLECTS in BETA band during all tasks.

| | | Region Pair | Verb Generation | | Repetition | | Resting | |
|---|---|---|---|---|---|---|---|---|
| | | | Estimate (95% CI) | p-value | Estimate (95% CI) | p-value | Estimate (95% CI) | p-value |
| Motor to Frontal | Intra-hemispheric | LMotor — LFront | *0.05 (-0.009, 0.1)* | *0.10* | *0.04 (-0.02, 0.09)* | *0.16* | *0.04 (-0.02, 0.09)* | *0.16* |
| | | RMotor — RFront | *-0.008 (-0.06, 0.04)* | *0.75* | *0.02 (-0.04, 0.09)* | *0.49* | *0.02 (-0.04, 0.09)* | *0.52* |
| | Inter-hemispheric | LMotor — RFront | *0.06 (-0.001, 0.1)* | *0.06* | *0.06 (-0.02, 0.14)* | *0.12* | *-0.003 (-0.06, 0.06)* | *0.93* |
| | | RMotor — LFront | *0.05 (-0.007, 0.1)* | *0.09* | *0.06 (-0.001, 0.1)* | *0.05* | *0.03 (-0.04, 0.09)* | *0.41* |
| Motor to Temporal | Intra-hemispheric | LMotor — LTemp | *0.02 (-0.04, 0.08)* | *0.55* | *0.02 (-0.05, 0.09)* | *0.60* | *0.02 (-0.03, 0.08)* | *0.43* |
| | | RMotor — RTemp | *0.03 (-0.03, 0.1)* | *0.35* | *0.06 (-0.01, 0.1)* | *0.11* | *0.006 (-0.05, 0.06)* | *0.84* |
| | Inter-hemispheric | LMotor — RTemp | *0.06 (0.009, 0.1)* | *0.02* | *0.05 (-0.01, 0.1)* | *0.11* | *0.03 (-0.02, 0.07)* | *0.22* |
| | | RMotor — LTemp | *0.04 (-0.01, 0.08)* | *0.15* | *0.05 (-0.02, 0.1)* | *0.14* | *0.03 (-0.01, 0.07)* | *0.18* |
| Frontal to Temporal | Intra-hemispheric | LFront — LTemp | *0.04 (-0.03, 0.1)* | *0.26* | *0.02 (-0.07, 0.1)* | *0.70* | *0.01 (-0.06, 0.09)* | *0.73* |
| | | RFront — RTemp | *-0.004 (-0.09, 0.08)* | *0.92* | *0.02 (-0.07, 0.1)* | *0.64* | *-0.02 (-0.1, 0.05)* | *0.54* |
| | Inter-hemispheric | RFront — LTemp | *0.03 (-0.03, 0.09)* | *0.39* | *0.02 (-0.06, 0.1)* | *0.59* | *-0.01 (-0.08, 0.05)* | *0.70* |
| | | LFront — RTemp | *0.01 (-0.06, 0.08)* | *0.79* | *0.03 (-0.05, 0.1)* | *0.45* | *0.004 (-0.06, 0.06)* | *0.89* |

P-values meeting threshold ($p<0.0042$) are in bold. LFront: Left Inferior Frontal; LTemp: Left Superior Temporal; LMotor: Left Motor; RFront: Right Inferior Frontal; RTemp: Right Superior Temporal; RMotor: Right Motor.

**Supplemental Table 4.** Group differences in connectivity for Children with SeLECTS in GAMMA band during all tasks.

| | | Region Pair | Verb Generation | | Repetition | | Resting | |
|---|---|---|---|---|---|---|---|---|
| | | | Estimate (95% CI) | p-value | Estimate (95% CI) | p-value | Estimate (95% CI) | p-value |
| Motor to Frontal | Intra-hemispheric | LMotor — LFront | *-0.01 (-0.09, 0.06)* | *0.70* | *0.01 (-0.06, 0.09)* | *0.72* | *0.03 (-0.04, 0.09)* | *0.40* |
| | | RMotor — RFront | *-0.01 (-0.1, 0.08)* | *0.83* | *0.01 (-0.08, 0.1)* | *0.82* | *0.03 (-0.06, 0.1)* | *0.52* |
| | Inter-hemispheric | LMotor — RFront | *0.04 (-0.01, 0.1)* | *0.12* | *0.05 (-0.01, 0.1)* | *0.11* | *0.03 (-0.01, 0.08)* | *0.14* |
| | | RMotor — LFront | *0.01 (-0.04, 0.07)* | *0.66* | *0.05 (-0.02, 0.1)* | *0.13* | *0.01 (-0.05, 0.08)* | *0.69* |
| Motor to Temporal | Intra-hemispheric | LMotor — LTemp | *-0.03 (-0.1, 0.05)* | *0.49* | *-0.002 (-0.08, 0.08)* | *0.97* | *0.003 (-0.05, 0.06)* | *0.90* |
| | | RMotor — RTemp | *-0.03 (-0.1, 0.04)* | *0.36* | *-0.004 (-0.09, 0.08)* | *0.93* | *-0.005 (-0.08, 0.07)* | *0.90* |
| | Inter-hemispheric | LMotor — RTemp | *0.03 (-0.03, 0.09)* | *0.31* | *0.02 (-0.04, 0.08)* | *0.48* | *0.03 (-0.02, 0.09)* | *0.18* |
| | | RMotor — LTemp | *0.01 (-0.05, 0.08)* | *0.72* | *0.02 (-0.05, 0.08)* | *0.62* | *0.01 (-0.03, 0.05)* | *0.64* |
| Frontal to Temporal | Intra-hemispheric | LFront — LTemp | *-0.002 (-0.09, 0.09)* | *0.96* | *-0.02 (-0.1, 0.08)* | *0.73* | *0.006 (-0.06, 0.07)* | *0.87* |
| | | RFront — RTemp | *-0.03 (-0.1, 0.06)* | *0.55* | *0.002 (-0.1, 0.1)* | *0.96* | *0.009 (-0.09, 0.1)* | *0.86* |
| | Inter-hemispheric | RFront — LTemp | *0.01 (-0.04, 0.07)* | *0.71* | *0.03 (-0.05, 0.1)* | *0.43* | *0.02 (-0.06, 0.09)* | *0.63* |
| | | LFront — RTemp | *0.01 (-0.04, 0.07)* | *0.61* | *0.05 (-0.02, 0.1)* | *0.18* | *-0.0005 (-0.05, 0.05)* | *0.98* |

P-values meeting threshold ($p<0.0042$) are in bold. LFront: Left Inferior Frontal; LTemp: Left Superior Temporal; LMotor: Left Motor; RFront: Right Inferior Frontal; RTemp: Right Superior Temporal; RMotor: Right Motor.

**Supplementary Material & Figure 1. Changes in wPLI connectivity estimated from sliding time window analysis.**

**Rationale:** Our primary analyses average over a 3 second epoch, which may miss important temporally specific changes in connectivity as language processing occurs within hundreds of milliseconds. Phase-based connectivity measures require 3-5 cycles to be reliable, and hence theta connectivity requires ~1 second epochs. Here, we performed a sliding scale analysis to test whether our inferences about group or task connectivity differences changed when using more temporally precise epochs.

**Methods:** We re-segmented data into shorter time windows, including the time window immediately before noun onset (-1 to 0 seconds), immediately after noun presentation (0 to 1 second), 0.5 to 1.5 seconds after noun presentation, and 1 to 2 seconds after noun presentation. We fit a GEE model with theta wPLI connectivity as the dependent variable and group (SeLECTS/controls), time (-1 to 0 second; 0 to 1 second; 0.5 to 1.5 seconds; 1 to 2 seconds), and the group by time interaction as independent variables, with adjustment for age and sex; we performed analyses for the verb generation task. We ran separate models for each of the 12 region-to-region connectivity pairs, and considered $p < 0.0042$ significant as per Bonferroni correction.

**Results:**

**Supplemental Figure 1.** Time period group differences in connectivity for Children with SeLECTS and Controls during the verb generation task (significant group differences in red p-value).

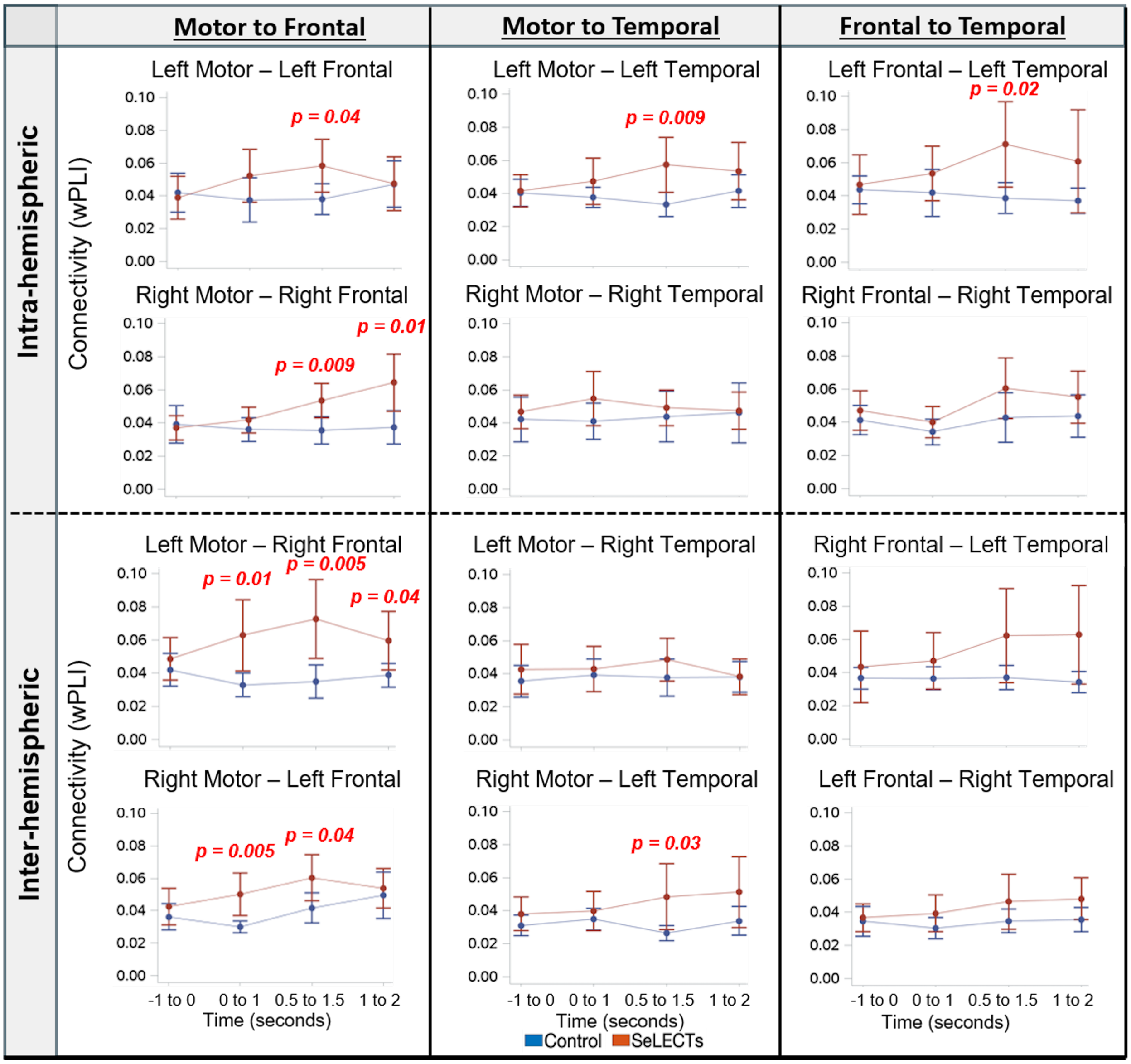


Plots represent estimated marginal mean connectivity with 95% confidence intervals for each group. P-values meeting threshold ($p<0.0042$) noted and color coded to group (red=SeLECTS; blue=controls). Left column: Motor to Inferior Frontal connectivity; Middle column: Motor to Superior Temporal connectivity; Right column: Inferior Frontal to Superior Temporal connectivity.

**Conclusion:** As our results were consistent with analyses using longer 3 second epochs, our inferences about the previously reported group differences in connectivity do not change significantly with enhanced temporal precision.

**Supplemental Table 5.** Subject Demographics by Medication Usage

| **Demographics** | **Group** | | | |
|---|---|---|---|---|
| | SeLECTS-ASM (n=14) | SeLECTS+ASM (n=17) | Controls (n=32) | p-value |
| Age (years) mean, SD | *10.2 +/- 2.0* | *9.3 +/- 1.9* | *9.1 +/- 2.0* | *0.20* |
| Sex (male), n (%) | *10 (71)* | *12 (71)* | *18 (56)* | *0.48* |
| Edinburgh Handedness Inventory* | *0.83 +/- 0.09* | *0.80+/- 0.17* | *0.77 +/- 0.17* | *0.45* |
| Hollingshead SES Index | *56.3 +/- 9.8* | *53.2 +/- 12.4* | *60.3 +/- 4.4* | *0.03* |
| **Neuropsychological Testing** | SeLECTS-ASM (n=12) | SeLECTS+ASM (n=16) | Controls (n=30) | p-value |
| WASI-II IQ | *105.7 +/- 13.6* | *97+/- 14.0* | *115.2 +/- 14.8* | *0.0006* |
| CTOPP-2 Phonological Awareness Score | *107.6 +/- 13.7* | *98.4 +/- 13.4* | *113.4 +/- 13.4* | *0.003* |

*Edinburgh Handedness Inventory (+1 = strongly right-handed and -1 = left-handed); ADHD: Attention Deficit Hyperactivity Disorder; CTOPP-2: Comprehensive Test of Phonological Processing-2nd Edition; SeLECTS-ASM: SeLECTS children not taking a daily antiseizure medication; SeLECTS+ASM: SeLECTS children taking a daily antiseizure medication; SES: Socioeconomic Status; WASI-II IQ: Wechsler Abbreviated Scale of Intelligence-2nd Edition Intelligence Quotient.

**Supplemental Table 6.** Group differences in connectivity for Children with SeLECTS off ASM and Controls during all tasks.

| | | Region Pair | Verb generation | | Repetition | | Resting | |
|---|---|---|---|---|---|---|---|---|
| | | | Estimate (95% CI) | p-value | Estimate (95% CI) | p-value | Estimate (95% CI) | p-value |
| Motor to Frontal | Intra-hemispheric | LMotor — LFront | *-0.006 (-0.05, 0.03)* | *0.77* | *0.0009 (-0.03, 0.04)* | *0.96* | *-0.005 (-0.05, 0.04)* | *0.86* |
| | | RMotor — RFront | *0.03 (-0.001, 0.06)* | *0.06* | *0.03 (-0.007, 0.06)* | *0.12* | *0.04 (-0.003, 0.09)* | *0.07* |
| | Inter-hemispheric | LMotor — RFront | *0.04 (-0.03, 0.11)* | *0.24* | *0.03 (-0.02, 0.08)* | *0.18* | *0.01 (-0.03, 0.06)* | *0.61* |
| | | RMotor — LFront | *0.007 (-0.03, 0.04)* | *0.68* | *0.009 (-0.02, 0.03)* | *0.48* | *0.01 (-0.03, 0.05)* | *0.52* |
| Motor to Temporal | Intra-hemispheric | LMotor — LTemp | *0.01 (-0.02, 0.05)* | *0.47* | *-0.01 (-0.05, 0.02)* | *0.53* | *0.03 (-0.01, 0.07)* | *0.18* |
| | | RMotor — RTemp | *0.03 (-0.02, 0.09)* | *0.26* | *0.03 (-0.02, 0.07)* | *0.22* | *0.02 (-0.03, 0.08)* | *0.38* |
| | Inter-hemispheric | LMotor — RTemp | *-0.001 (-0.03, 0.02)* | *0.91* | *-0.002 (-0.04, 0.03)* | *0.90* | *-0.02 (-0.04, 0.01)* | *0.30* |
| | | RMotor — LTemp | *0.005 (-0.02, 0.03)* | *0.63* | *-0.001 (-0.02, 0.02)* | *0.93* | *0.02 (-0.002, 0.04)* | *0.08* |
| Frontal to Temporal | Intra-hemispheric | LFront — LTemp | *0.006 (-0.03, 0.04)* | *0.73* | *-0.02 (-0.06, 0.02)* | *0.41* | *0.01 (-0.04, 0.07)* | *0.65* |
| | | RFront — RTemp | *-0.008 (-0.04, 0.03)* | *0.63* | *-0.04 (-0.08, 0.007)* | *0.10* | *-0.03 (-0.08, 0.02)* | *0.24* |
| | Inter-hemispheric | RFront — LTemp | *0.008 (-0.03, 0.05)* | *0.70* | *-0.005 (-0.04, 0.03)* | *0.78* | *0.03 (-0.01, 0.07)* | *0.16* |
| | | LFront — RTemp | *-0.005 (-0.04, 0.03)* | *0.78* | *-0.01 (-0.04, 0.01)* | *0.24* | *-0.02 (-0.05, 0.003)* | *0.08* |

P-values meeting threshold ($p<0.0042$) are in bold. LFront: Left Inferior Frontal; LTemp: Left Superior Temporal; LMotor: Left Motor; RFront: Right Inferior Frontal; RTemp: Right Superior Temporal; RMotor: Right Motor.

**Supplemental Table 7.** Group differences in connectivity for Children with SeLECTS on ASM and Controls during all tasks.

| | | Region Pair | Verb generation | | Repetition | | Resting | |
|---|---|---|---|---|---|---|---|---|
| | | | Estimate (95% CI) | p-value | Estimate (95% CI) | p-value | Estimate (95% CI) | p-value |
| Motor to Frontal | Intra-hemispheric | LMotor — LFront | *0.08 (0.01, 0.16)* | *0.02* | *0.06 (0.004, 0.11)* | *0.04* | *0.08 (0.01, 0.14)* | *0.02* |
| | | RMotor — RFront | *0.08 (0.04, 0,12)* | ***0.0001*** | *0.08 (0.04, 0.11)* | ***0.0001*** | *0.09 (0.02, 0.17)* | *0.02* |
| | Inter-hemispheric | LMotor — RFront | *0.08 (0.03, 0.12)* | ***0.001*** | *0.09 (0.04, 0.13)* | ***0.0003*** | *0.03 (-0.009, 0.08)* | *0.12* |
| | | RMotor — LFront | *0.06 (0.006, 0.11)* | *0.03* | *0.05 (0.02, 0.08)* | ***0.0004*** | *0.06 (0.02, 0.11)* | *0.009* |
| Motor to Temporal | Intra-hemispheric | LMotor — LTemp | *0.07 (0.01, 0.12)* | *0.02* | *0.03 (-0.01, 0.07)* | *0.16* | *0.05 (0.01, 0.08)* | *0.008* |
| | | RMotor—RTemp | *0.02 (-0.03, 0.07)* | *0.42* | *0.03 (-0.006, 0.07)* | *0.10* | *0.03 (-0.01, 0.08)* | *0.13* |
| | Inter-hemispheric | LMotor — RTemp | *0.03 (-0.01, 0.07)* | *0.16* | *0.03 (-0.01, 0.07)* | *0.15* | *0.01 (-0.02, 0.05)* | *0.54* |
| | | RMotor — LTemp | *0.07 (0.01, 0.12)* | *0.01* | *0.03 (-0.02, 0.08)* | *0.25* | *0.04 (0.01, 0.07)* | ***0.003*** |
| Frontal to Temporal | Intra-hemispheric | LFront — LTemp | *0.13 (0.05, 0.2)* | ***0.0009*** | *0.08 (0.0003, 0.16)* | *0.05* | *0.05 (-0.01, 0.12)* | *0.11* |
| | | RFront — RTemp | *0.03 (-0.02, 0.08)* | *0.21* | *0.06 (-0.009, 0.13)* | *0.09* | *0.03 (-0.04, 0.10)* | *0.35* |
| | Inter-hemispheric | RFront — LTemp | *0.08 (0.02, 0.14)* | *0.01* | *0.05 (-0.01, 0.11)* | *0.13* | *0.03 (-0.0008, 0.07)* | *0.06* |
| | | LFront — RTemp | *0.05 (-0.005, 0.11)* | *0.07* | *0.04 (0.007, 0.08)* | *0.02* | *0.04 (-0.006, 0.08)* | *0.10* |

P-values meeting threshold (p<0.0042) are in bold. LFront: Left Inferior Frontal; LTemp: Left Superior Temporal; LMotor: Left Motor; RFront: Right Inferior Frontal; RTemp: Right Superior Temporal; RMotor: Right Motor.

**Supplemental Table 8.** Group differences in connectivity for Children with SeLECTS on and off ASM during all tasks.

| | | Region Pair | Verb generation | | Repetition | | Resting | |
|---|---|---|---|---|---|---|---|---|
| | | | Estimate (95% CI) | p-value | Estimate (95% CI) | p-value | Estimate (95% CI) | p-value |
| Motor to Frontal | Intra-hemispheric | LMotor — LFront | *0.09 (0.02, 0.16)* | *0.01* | *0.06 (0.004, 0.11)* | *0.04* | *0.08 (0.01, 0.15)* | *0.02* |
| | | RMotor — RFront | *0.05 (0.009, 0.10)* | *0.38* | *0.05 (0.008, 0.1)* | *0.02* | *0.05 (-0.04, 0.14)* | *0.25* |
| | Inter-hemispheric | LMotor — RFront | *0.04 (-0.04, 0.12)* | *0.38* | *0.05 (-0.01, 0.12)* | *0.12* | *0.02 (-0.03, 0.08)* | *0.39* |
| | | RMotor — LFront | *0.05 (-0.005, 0.10)* | *0.07* | *0.04 (0.01, 0.08)* | *0.01* | *0.05 (-0.005, 0.11)* | *0.07* |
| Motor to Temporal | Intra-hemispheric | LMotor — LTemp | *0.05 (-0.002, 0.11)* | *0.06* | *0.04 (-0.006, 0.09)* | *0.09* | *0.02 (-0.02, 0.07)* | *0.35* |
| | | RMotor — RTemp | *-0.01 (-0.07, 0.05)* | *0.71* | *0.005 (-0.03, 0.04)* | *0.80* | *0.01 (-0.05, 0.07)* | *0.71* |
| | Inter-hemispheric | LMotor — RTemp | *0.03 (-0.01, 0.08)* | *0.18* | *0.03 (-0.01, 0.08)* | *0.16* | *0.03 (-0.006, 0.06)* | *0.11* |
| | | RMotor — LTemp | *0.06 (0.003, 0.12)* | *0.04* | *0.03 (-0.03, 0.08)* | *0.30* | *0.03 (-0.007, 0.06)* | *0.12* |
| Frontal to Temporal | Intra-hemispheric | LFront — LTemp | *0.12 (0.04, 0.20)* | ***0.003*** | *0.1 (0.02, 0.18)* | *0.02* | *0.04 (-0.04, 0.12)* | *0.29* |
| | | RFront — RTemp | *0.04 (-0.02, 0.10)* | *0.17* | *0.1 (0.02, 0.17)* | *0.01* | *0.06 (-0.01, 0.13)* | *0.09* |
| | Inter-hemispheric | RFront — LTemp | *0.07 (-0.001, 0.15)* | *0.05* | *0.05 (-0.02, 0.12)* | *0.16* | *0.004 (-0.05, 0.06)* | *0.86* |
| | | LFront — RTemp | *0.06 (-0.008, 0.12)* | *0.09* | *0.06 (0.02, 0.10)* | ***0.003*** | *0.06 (0.02, 0.10)* | *0.0043* |

P-values meeting threshold (p<0.0042) are in bold. LFront: Left Inferior Frontal; LTemp: Left Superior Temporal; LMotor: Left Motor; RFront: Right Inferior Frontal; RTemp: Right Superior Temporal; RMotor: Right Motor.

**Supplemental Methods & Table 9.** Epilepsy severity for children with SeLECTS.

**Assessment of Epilepsy Severity:** There is not a validated scale of epilepsy severity for SeLECTS. The Global assessment of the severity of epilepsy (GASE) is a scale validated for assessing epilepsy severity in children that is filled out by the patient's treating physician. We chose questions from this assessment that were relevant to SeLECTS and could be extracted from the medical record to compare epilepsy between children with SeLECTS taking vs. not taking daily prophylactic ASMs. We compared groups using the t-test and chi-squared test for continuous and categorical values.

| | SeLECTS - ASM (n=14) | SeLECTS + ASM (n=17) | test statistic | p-value |
|---|---|---|---|---|
| Number of Lifetime Seizures | *7.3 +/- 10.3* | *12.9 +/- 12.8* | *1.3* | *0.21* |
| Hx Seizures > 5min, n (%) | *2 (14)* | *2 (12)* | *0.11* | *0.74* |
| Seizure state | | | | |
| *Asleep Only* | *9 (64)* | *6 (35)* | *1.6* | *0.21* |
| *Awake & Asleep* | *5 (36)* | *11 (65)* | *1.6* | *0.21* |
| Hx Seizures with Secondary Generalization, n (%) | *10 (71)* | *12 (71)* | *0.12* | *0.73* |

ASM: Antiseizure medication; Hx: history; SeLECTS-ASM: SeLECTS children not taking a daily antiseizure medication; SeLECTS+ASM: SeLECTS children taking a daily antiseizure medication.

**Conclusion:** Children with SeLECTS taking a daily antiseizure medication had a higher number of lifetime seizures, more seizures when both awake and asleep, and a history of seizures with more secondary generalization, though none of these differences were significant between groups.

**Supplemental Figure 2.** Association between Clinical Language Performance and Connectivity in Children with SeLECTS and Controls during the Repetition task.

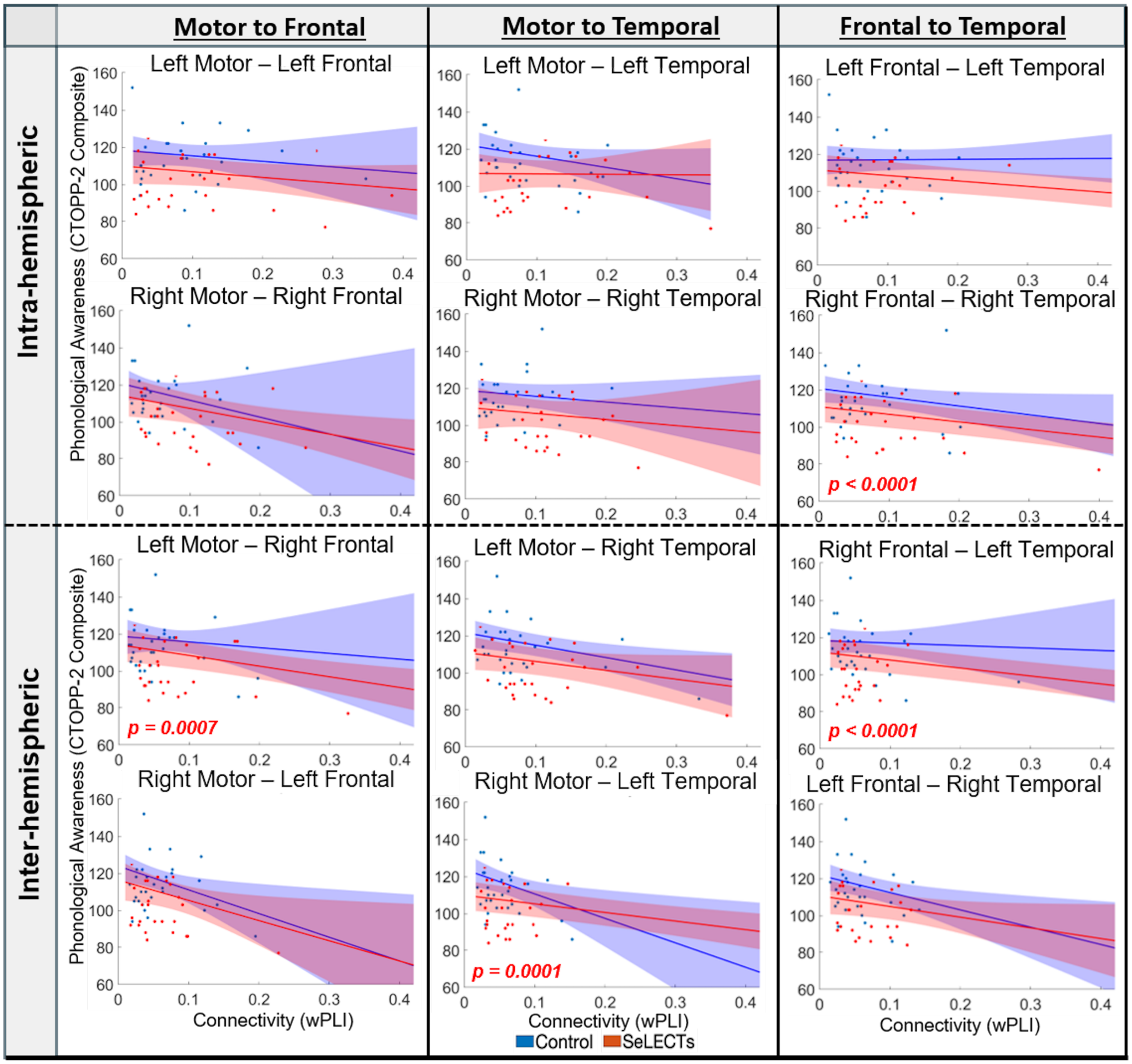


Scatter plots of mean connectivity vs. CTOPP-2 Composite scores for each subject and estimated marginal fit lines with 95% confidence intervals. P-values meeting threshold ($p<0.0042$) noted and color coded to group (red=SeLECTS; blue=controls). Left column: Motor to Inferior Frontal; Middle column: Motor to Superior Temporal; Right column: Inferior Frontal to Superior Temporal.

**Supplemental Figure 3.** Association between Clinical Language Performance and Connectivity in Children with SeLECTS and Controls during the Resting task.

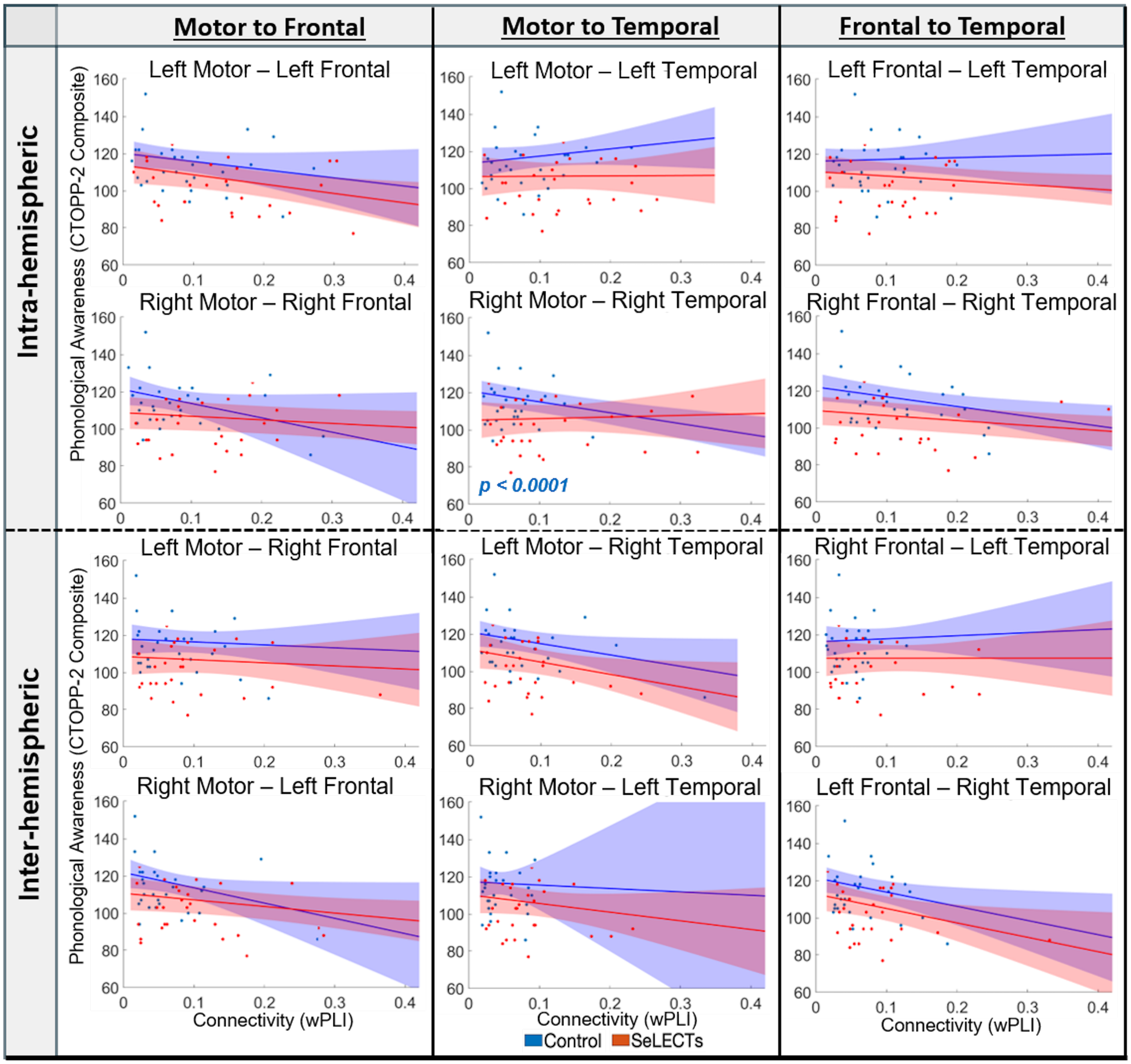


Scatter plots of mean connectivity vs. CTOPP-2 Composite scores for each subject and estimated marginal fit lines with 95% confidence intervals. P-values meeting threshold ($p<0.0042$) noted and color coded to group (red=SeLECTS; blue=controls). Left column: Motor to Inferior Frontal; Middle column: Motor to Superior Temporal; Right column: Inferior Frontal to Superior Temporal.

**Supplemental Table 10.** Association between Clinical Language Performance and Connectivity in Children with SeLECTS adjusting for age, sex, socioeconomic status, IQ, and Handedness.

| | | Region Pair | Verb generation | | Repetition | | Resting | |
|---|---|---|---|---|---|---|---|---|
| | | | Estimate (95% CI) | p-value | Estimate (95% CI) | p-value | Estimate (95% CI) | p-value |
| Motor to Frontal | Intra-hemispheric | LMotor — LFront | *-23.5 (-60.0, 12.9)* | *0.21* | *-22.4 (-65.2, 20.4)* | *0.3* | *-41.9 (-80.0, -3.8)* | *0.03* |
| | | RMotor — RFront | *-72.8 (-137.8, -7.7)* | *0.03* | *-54.1 (-138.9, 30.7)* | *0.21* | *-21.0 (-49.1, 7.0)* | *0.14* |
| | Inter-hemispheric | LMotor — RFront | *-55.3 (-103.2, -7.3)* | *0.02* | *-49.5 (-107.8, 8.9)* | *0.1* | *14.2 (-43.4, 71.8)* | *0.63* |
| | | RMotor — LFront | *-39.2 (-98.2, 19.8)* | *0.19* | *-61.2 (-173.1, 50.8)* | *0.28* | *-17.9 (-54.7, 19.0)* | *0.34* |
| Motor to Temporal | Intra-hemispheric | LMotor — LTemp | *0.74 (-64.2, 65.7)* | *0.98* | *2.7 (-80.7, 86.2)* | *0.95* | *33.9 (-33.2, 100.9)* | *0.32* |
| | | RMotor — RTemp | *-15.8 (-92.2, 60.7)* | *0.69* | *-39.1 (-142.8, 64.6)* | *0.46* | *28.8 (-21.1, 78.7)* | *0.26* |
| | Inter-hemispheric | LMotor — RTemp | *-64.2 (-101.6, -26.9)* | ***0.0008*** | *-47.5 (-105.9, 11.0)* | *0.11* | *1.0 (-61.7, 63.8)* | *0.97* |
| | | RMotor — LTemp | *-62.0 (-84.4, -39.6)* | ***0.0001*** | *-52.7 (-77.6, -27.8)* | ***0.0001*** | *77.7 (-22.1, 177.4)* | *0.13* |
| Frontal to Temporal | Intra-hemispheric | LFront — LTemp | *-29.4 (-59.8, 1.0)* | *0.06* | *-27.5 (-54.8, -0.29)* | *0.05* | *-9.9 (-39.6, 19.8)* | *0.51* |
| | | RFront — RTemp | *-62.2 (-110.0, -14.5)* | *0.01* | *-44.2 (-62.7, -25.6)* | ***0.0001*** | *-11.6 (-34.6, 11.4)* | *0.32* |
| | Inter-hemispheric | RFront — LTemp | *-47.1 (-70.2, -24.0)* | ***0.0001*** | *-39.6 (-58.7, -20.5)* | ***0.0001*** | *45.2 (-7.7, 98.1)* | *0.09* |
| | | LFront — RTemp | *-48.5 (-72.9, -24.1)* | ***0.0001*** | *-62.8 (-128.4, 2.9)* | *0.06* | *-45.9 (-111.2, 19.3)* | *0.17* |

P-values meeting threshold ($p<0.0042$) are in bold. LFront: Left Inferior Frontal; LTemp: Left Superior Temporal; LMotor: Left Motor; RFront: Right Inferior Frontal; RTemp: Right Superior Temporal; RMotor: Right Motor.

**Supplemental Table 11.** Association between Clinical Language Performance and Connectivity in Controls adjusting for age, sex, socioeconomic status, IQ, and Handedness.

| | | Region Pair | Verb generation | | Repetition | | Resting | |
|---|---|---|---|---|---|---|---|---|
| | | | Estimate (95% CI) | p-value | Estimate (95% CI) | p-value | Estimate (95% CI) | p-value |
| Motor to Frontal | Intra-hemispheric | LMotor — LFront | *-58.8 (-105.7, -11.9)* | *0.01* | *-38.6 (-100.1, 22.9)* | *0.22* | *-46.6 (-93.7, 0.5)* | *0.05* |
| | | RMotor — RFront | *-70.1 (-157.9, 17.7)* | *0.12* | *-87.3 (-216.4, 41.8)* | *0.19* | *-84.6 (-152.4, -16.8)* | *0.01* |
| | Inter-hemispheric | LMotor — RFront | *-39.0 (-132.0, 54.1)* | *0.41* | *6.5 (-78.9, 91.9)* | *0.88* | *-31.5 (-84.7, 21.7)* | *0.25* |
| | | RMotor — LFront | *-24.6 (-105.8, 56.6)* | *0.55* | *-88.2 (-199.9, 23.4)* | *0.12* | *-81.8 (-149.5, -14.1)* | *0.02* |
| Motor to Temporal | Intra-hemispheric | LMotor — LTemp | *-62.4 (-165.1, 40.4)* | *0.23* | *-28.9 (-87.2, 29.3)* | *0.33* | *37.4 (-15.5, 90.3)* | *0.17* |
| | | RMotor — RTemp | *-28.1 (-70.7, 14.4)* | *0.19* | *-24.3 (-75.9, 27.3)* | *0.36* | *-55.6 (-82.2, -29.0)* | ***0.0001*** |
| | Inter-hemispheric | LMotor — RTemp | *-13.4 (-142.1, 115.2)* | *0.84* | *-64.0 (-104.6, -23.4)* | ***0.002*** | *-67.8 (-106.5, -29.0)* | ***0.0006*** |
| | | RMotor — LTemp | *-104.8 (-175.0, -34.6)* | ***0.003*** | *-123.0 (-215.5, -30.4)* | *0.009* | *-84.4 (-278.1, 109.2)* | *0.39* |
| Frontal to Temporal | Intra-hemispheric | LFront — LTemp | *-2.0 (-84.7, 80.7)* | *0.96* | *-1.1 (-46.2, 43.9)* | *0.96* | *-3.5 (-51.7, 44.7)* | *0.89* |
| | | RFront — RTemp | *-32.5 (-91.4, 26.4)* | *0.28* | *-34.8 (-78.8, 9.2)* | *0.12* | *-45.8 (-77.7, -14.0)* | *0.005* |
| | Inter-hemispheric | RFront — LTemp | *-14.3 (-123.4, 94.8)* | *0.80* | *12.1 (-71.4, 95.5)* | *0.78* | *-1.2 (-82.2, 79.8)* | *0.98* |
| | | LFront — RTemp | *-57.1 (-124.5, 10.3)* | *0.10* | *-75.5 (-142.9, -8.1)* | *0.03* | *-79.8 (-127.5, -32.1)* | ***0.001*** |

P-values meeting threshold ($p<0.0042$) are in bold. LFront: Left Inferior Frontal; LTemp: Left Superior Temporal; LMotor: Left Motor; RFront: Right Inferior Frontal; RTemp: Right Superior Temporal; RMotor: Right Motor.
